# Large- and multi-scale networks in the rodent brain during novelty exploration

**DOI:** 10.1101/2020.12.08.416248

**Authors:** Michael X Cohen, Bernhard Englitz, Arthur S C França

## Abstract

Neural activity is coordinated across multiple spatial and temporal scales, and these patterns of coordination are implicated in both healthy and impaired cognitive operations. However, empirical cross-scale investigations are relatively infrequent, due to limited data availability and to the difficulty of analyzing rich multivariate datasets. Here we applied frequency-resolved multivariate source-separation analyses to characterize a large-scale dataset comprising spiking and local field potential activity recorded simultaneously in three brain regions (prefrontal cortex, parietal cortex, hippocampus) in freely-moving mice. We identified a constellation of multidimensional, inter-regional networks across a range of frequencies (2-200 Hz). These networks were reproducible within animals across different recording sessions, but varied across different animals, suggesting individual variability in network architecture. The theta band (~4-10 Hz) networks had several prominent features, including roughly equal contribution from all regions and strong inter-network synchronization. Overall, these findings demonstrate a multidimensional landscape of large-scale functional activations of cortical networks operating across multiple spatial, spectral, and temporal scales during open-field exploration.

**Significance statement:** Neural activity is synchronized over space, time, and frequency. To characterize the dynamics of large-scale networks spanning multiple brain regions, we recorded data from the prefrontal cortex, parietal cortex, and hippocampus in awake behaving mice, and pooled data from spiking activity and local field potentials into one data matrix. Frequency-specific multivariate decomposition methods revealed a cornucopia of neural networks defined by coherent spatiotemporal patterns over time. These findings reveal a rich, dynamic, and multivariate landscape of large-scale neural activity patterns during foraging behavior.

## Introduction

Neural activity is coordinated across multiple spatial and temporal scales, ranging from spike-timing correlations across pairs of neurons (Gray et al., 1989) to resting-state fMRI networks (Gusnard et al., 2001), and from ultra-fast 600 Hz omega oscillations in primary sensory cortex (Timofeev and Bazhenov, 2005) to infra-slow fluctuations linked to 0.05 Hz oscillations in the gastric system (Richter et al., 2017). Coordinated activity is thought to allow for neural circuits to maximize communication efficiency, multiplex information, flexibly route information flow, and functionally bind cell assemblies (Jensen and Mazaheri, 2010; Singer, 2009; Wang, 2010).

However, most neuroscience investigations are limited to a single spatial scale, and cross-scale investigations are often limited to univariate or bivariate analyses, e.g., coherence between action potentials from one neuron with the LFP recorded on the same electrode (Whittingstall K Logothetis, 2009). This mass-univariate approach has been crucial to the development of neuroscience, for example, understanding of computational principles such as neural tuning (Carandini, 2005; Hebart and Baker, 2018; Hubel and Wiesel, 1959). However, it may obscure spatiotemporal patterns embedded across populations of neurons within and across brain regions (Cunningham and Yu, 2014; Kriegeskorte and Kievit, 2013; Ritchie et al., 2019; Williamson et al., 2019).

In contrast, multivariate data analysis methods have proven useful at identifying spatially distributed patterns that reflect lower-dimensional dynamics or that encode sensory representations or memories (Pang et al., 2016). Furthermore, correlational patterns may provide a “contextual activation” that shapes subsequent local computations (Alishbayli et al., 2019; Cohen and Kohn, 2011; Kohn et al., 2016; Priesemann et al., 2014).

In the present study, a recently developed set of multivariate methods (generalized eigendecomposition; (Cohen, 2017) enabled us to discover multi-scale, inter-regional functional networks during active behavior by combining data from multiunits and local field potentials. We found a salient, empirical grouping of the networks into a small number of frequency bands (average of 7). In each frequency bank, multiple subnetworks were both simultaneously and independently active. Some networks (e.g., in theta) were spatially distributed across the brain, while other networks (typically in higher frequencies) were more localized to one or two regions. Spiking activity contributed less systematically to brain-wide networks compared to LFP. The analyses revealed both idiosyncratic and reproducible network characteristics within- and across-animals, which suggests that the spatial organization of large-scale networks is subject to individual variability. Overall, our findings reveal a complex landscape of dynamic neural activity that spans multiple spatial, spectral, and temporal scales.

## Methods

### Data acquisition

Six male mice with Bl57/6jbackground (B6;129P2-Pvalbtm1(cr)Arbr/J or Ssttm2.1(cre)Zjh/J) between 4 and 5 months of age, weighing between 27g and 34g, were used in this study. All experiments were approved by the Dutch central commission for animal research (Centrale Commissie Dierproeven) and implemented according to approved work protocols from the local University Medical Centre animal welfare body (approval number 2016-0079).

Each animal was implanted with 32 electrodes (see Figure 1a) spread across the prefrontal cortex (16 electrodes), parietal cortex (8 electrodes), and hippocampus (8 electrodes). Inter-electrode distance was 250 μm and typical impedances were between 0.1 and 0.9 MOhm. Electrode design and surgeries are detailed elsewhere (van Hulten J. A. Cohen M. X, 2020). A metal reference screw was placed on the skull over the cerebellum. Offline, an average reference was computed for each brain region and subtracted from each electrode in the corresponding region.

**Figure 1.**
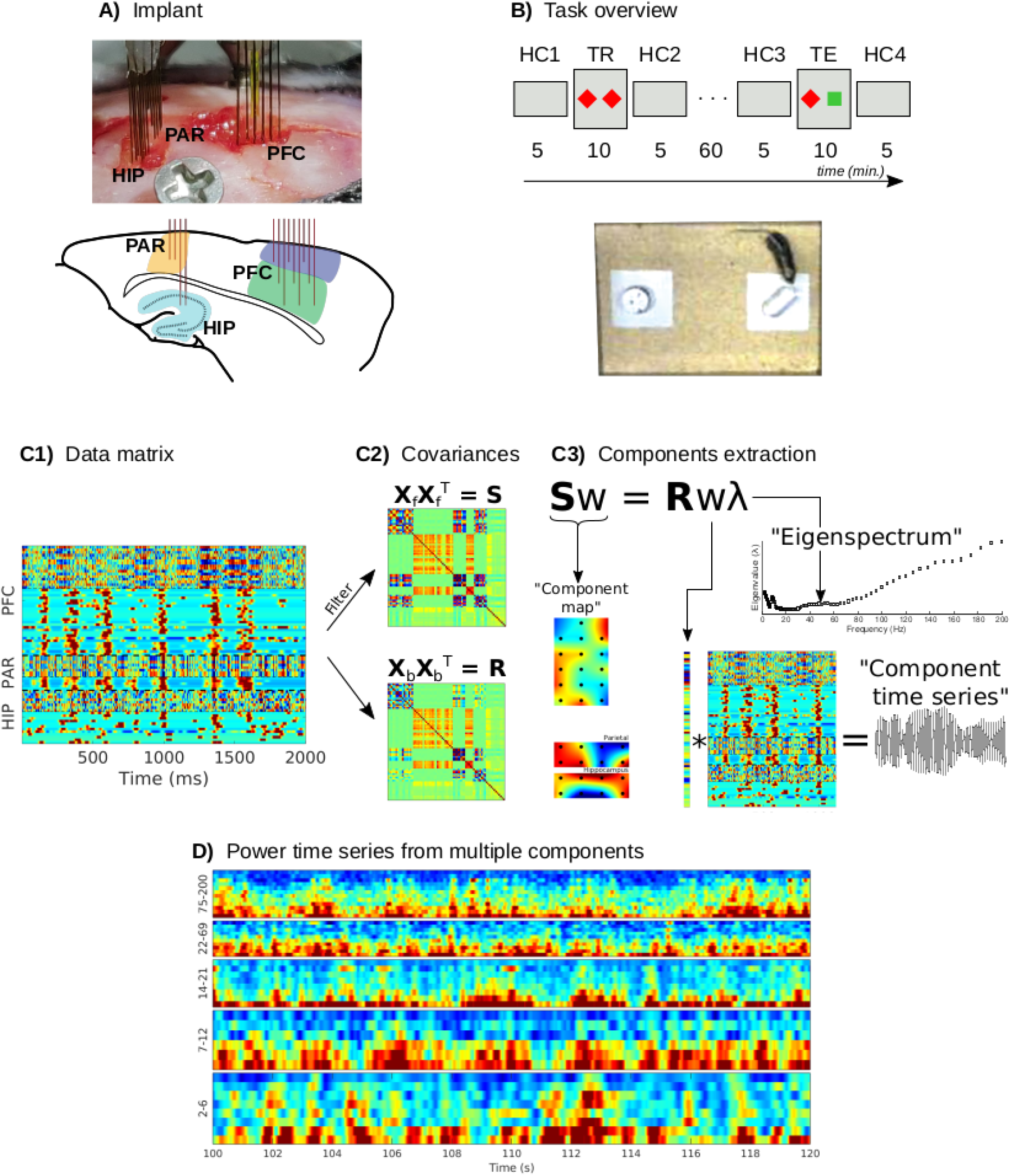
Overview of recording locations, task design, data analysis, and sample data. A) 32-channel custom-designed electrode array (HIP: hippocampus; PAR: parietal cortex; PFC: prefrontal cortex). The line drawing underneath illustrates the approximate locations of the electrodes on a sagittal slice. B) Task flow and timing (HC1-4: home cage sessions 1-4; TR: training; TE: testing). The red diamonds and green square indicate objects placed in the arena. The picture underneath is from a camera placed overhead. C1) A data matrix with combined LFP and multiunits (smoothed with a 30-ms FWHM Gaussian) from three different regions. C2) Data covariance matrices for the data snippet shown in C1, either narrowband-filtered (**S**) or broadband (**R**). A generalized eigendecomposition of these two matrices (panel C3) provides a set of eigenvectors (**w**) and corresponding eigenvalues (λ), from which three pieces of information are extracted: The component spatial map (the eigenvector multiplied by the covariance matrix), the component time series (the eigenvector multiplied by the data matrix), and the separability of narrowband vs. broadband activity (the eigenvalue for one frequency; the eigenvalues over frequencies creates an eigenspectrum). Illustrated here is one eigenvalue solution for one frequency; in practice, the number of solutions (**w**/λ pairs) corresponds to the number of data channels, and this entire procedure is repeated across a range of frequencies. D) Multiple components can be isolated from each frequency, with distinct temporal dynamics. Example component power time series are illustrated from 20 seconds of a recording; each row corresponds to a distinct component. Frequency groups are based on empirical frequency boundaries (described later) and components are sorted within each frequency band based on total component energy.

Animals were recorded in the sessions depicted in Figure 1b. The recording sessions alternated between their familiar home cage and an unfamiliar location that contained novel objects. In particular, each mouse went through the same succession of six experiment sessions: (1) Home cage recording of 5 minutes. (2) Training phase of 10 minutes, in which the animal was placed in an unfamiliar environment that contained two novel objects. (3) Home cage recording of 5 minutes. One hour then passed (in the home cage) with no recordings. (4) Home cage recording of 5 minutes. (5) Testing phase, in which the animal was returned to the unfamiliar environment that contained one object seen during the training phase and one novel object. (6) Home cage recording of 5 minutes. Mice were connected via electrode fibers to the data acquisition board via a cable that hung from top of the Faraday cage, but were otherwise unrestrained. There was no particular task or objective that was trained, nor were any rewards provided.

Mice tended to explore the objects for brief periods of time (hundreds of ms to seconds), whereas our data analysis approach utilized longer windows for temporal filtering and averaging to ensure high signal-to-noise quality. We therefore focused on possible state changes across the different task sessions as opposed to time-locking to the on/offsets of transient object exploration periods.

LFP data were down-sampled to 1000 Hz. Excessively noisy channels, determined based on visual inspection, were removed (0-4 per recording session; average of 1.2). Independent components analysis was run using the eeglab toolbox (Delorme and Makeig, 2004) and the jade algorithm (joint approximation diagonalization of eigen-matrices), which defines components by maximizing kurtosis (the 4th order statistical moment used to index non-Gaussianity) (Cardoso, 1999). Components clearly identifiable as non-neural origins were projected out of the data. Data from the first and last 10 seconds of each recording session were excluded from analyses.

Data and MATLAB analysis code will be made publically available upon acceptance via our institute’s data repository.

### Spike-sorting and multiunit extraction

The raw (30 kHz) voltage recordings were regional-average-referenced to eliminate possible volume-conduction artifacts, and were then filtered between 300 and 6000 Hz using a zero phase-shift FIR1 filter kernel. Spike-sorting was done for each electrode separately given the inter-electrode spacing of 250 μm, which makes it unlikely to observe the same neuron on multiple electrodes. Indeed, we did not find excessive correlations across units from different electrodes (see Figure 1-1 for an example between-unit correlation matrix).

Because our goal here was to obtain information about neural spiking activity as it related to the population and to LFP dynamics — rather than evaluating tuning properties of individual neurons — we chose an automatic spike-sorting approach that separated multiunits from noise or artifacts (Trautmann et al., 2019). We therefore term these signals “multiunit” to indicate that the resulting time series may reflect a mixture of action potentials from multiple neurons.

Multiunits were extracted via a general purpose spike-sorting suite (*autoSort*, available via our open code repository bitbucket.org/benglitz/controller-dnp/src/master/Access/SpikeSorting), implemented in MATLAB. Briefly, *autoSort* performs the following sequence of steps to achieve automatic and unbiased sorting of neural signals:

- Candidate spike waveforms ('spikes') were detected based on a negative threshold of 4 standard deviations of the background noise (estimated as 1.48 times the median absolute deviation, to avoid artifacts that inflate the standard deviation).
- Candidate spikes were then aligned to their minimum after the trigger and cut out within a window of [-0.7,1.2] ms relative to the alignment time.
- Principal components analysis was performed on a random subset of spikes (N_S_=5000 per recording) to estimate a projector to a 6 dimensional subspace that retained most of the variance in the data.
- Hierarchical clustering (based on *Ward* distance) with a set maximal number of clusters (N_C_=3) was performed on this representation, and all spikes beyond the N_S_-selection were assigned to these clusters on the basis of their Euclidean distance to the cluster centers.
- Clusters were then post-hoc automatically selected and fused on the basis of the shape and similarity between their average waveforms, i.e. (i) clusters were excluded if they had no significantly positive “hump” after the negative alignment peak, if they had a significantly positive peak before the negative alignment peak, or if the waveform was longer or larger than expected for an extracellular spike; and (ii) clusters were fused if the correlation and Euclidean distance between their average waveforms were above or below preset thresholds, respectively.

These steps and criteria led to an extraction of 0-2 multiunits per electrode. The average rate of spikes per second from all animals and recordings was 13.2 (standard deviation 5.9, minimum 0.07, maximum 51.6). A binary spike time series was constructed for each multiunit, and smoothed with a 30 ms full-width at half-maximum Gaussian to create a continuous signal. This continuous signal was entered into the data matrix as one channel (Figure 1C).

### Frequency-specific components using generalized eigendecomposition

We followed existing procedures for extracting multivariate components that have been detailed and validated in several previous publications (Cohen, 2017; de Cheveigné and Parra, 2014; Nikulin et al., 2011; Tomé, 2006). A brief overview is provided here.

The goal is to identify a spatial filter that provides a scalar weight for each data channel (LFP and multiunits) such that the weighted sum of narrowband-filtered channel time series is maximally different from the broadband channel time series. The method is based on data covariance matrices because they contain all pairwise linear relationships, making the method multivariate. As described below, two covariance matrices are compared, one matrix (R) based on the broadband (non-temporally filtered) data, and one matrix based on the narrowband filtered data (S).

Channel-by-channel covariance matrices were created by multiplying the mean-centered data matrices by their transpose. To increase covariance stability, we cut the continuous data into a series of non-overlapping 2 second segments, and computed the covariance matrix of each segment. The even-numbered epochs were used to create the S (signal) covariance matrix and the odd-numbered epochs were used to create the R (reference) covariance matrix. This was done to have non-identical data across the two matrices. After computing covariance matrices for each segment (there were around 70 segments in the home cage sessions and 140 segments in the training/testing sessions), the average covariance matrices S and R were computed across segments. Euclidean distance from each individual covariance matrix to the average was computed (this is equivalent to the Frobenius norm of the matrix difference), and any segments with a distance greater than three standard deviations from the average were excluded, and the final covariance matrix was re-computed without the outliers. On average, 0.85% of covariance matrices were excluded per analysis (range: 0 to 3%).

To create the spatial filter per frequency, we start from maximizing the Rayleigh quotient:

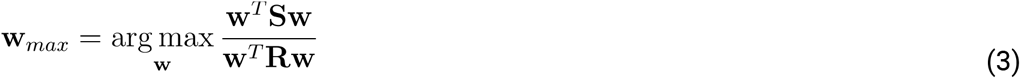

Where **S** and **R** are channel covariance matrices obtained from the narrowband filtered data and the broadband data, respectively (Figure 1C). One can think of equation 3 as a multivariate signal-to-noise ratio, and the goal is to find a channel vector **w** that maximizes this ratio. The solution comes from a generalized eigenvalue decomposition on the two matrices.

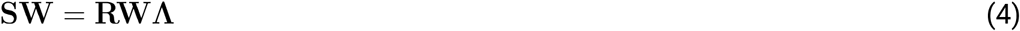

The diagonal matrix **Λ** contains the eigenvalues, each of which is the ratio of equation 3 for the corresponding column of **W**, which is a matrix in which the columns are the eigenvectors. Thus, we obtain *m* spatial filters for an *m*-channel dataset. The solutions are linearly independent from each other, though they are not constrained to be orthogonal as with PCA (this is because eigenvector orthogonality is guaranteed only for symmetric matrices, and **R**^-1^**S** is non-symmetric). Equation 4 is repeated for a range of temporal frequencies (see below), each using a different **S** matrix (the covariance matrix created from narrowband filtered data) with the same **R** matrix.

A small amount of shrinkage regularization (1%) was applied to the **R** matrix in order to improve the quality of the decomposition (Lotte and Guan, 2011). In our experience, 1% shrinkage has no appreciable effect on decompositions of clean, full-rank, and easily separable data, and considerably improves the decompositions of noisy or reduced-rank data. In equation 5 γ below, is the amount of shrinkage (0.01, corresponding to 1%), α is the average of all eigenvalues of **R**, and **I** is the identity matrix.

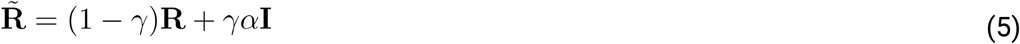

We refer to each spatial filter as a “component.” The component time series is obtained by multiplying **w** by the channels-by-time data matrix (this is how the eigenvector is a spatial filter). For all signals, any time series values exceeding four standard deviations from the mean of the time series were excluded. This reduced the possible impact of residual artifacts or edges influencing the results. The component map is obtained by multiplying **w** by the **S** covariance matrix (Haufe et al., 2014).

The entire procedure described above was repeated independently for each animal, experiment session, and filtering frequency. This allowed us to examine the reproducibility of the components both within and across animals.

Data were temporally narrowband filtered by convolution with a Morlet wavelet, defined here as a Gaussian in the frequency domain (Cohen, 2019). Extracted frequencies ranged from 2 Hz to 200 Hz in 100 logarithmically spaced steps. The full-width at half-maximum of the Gaussian varied from 2 to 5 Hz with increasing frequency. The multiunit channels were not narrowband filtered (they were already smoothed with a 30-ms Gaussian). Any large-scale spike-field coherence patterns would manifest as cross-channel terms in the frequency-specific covariance matrices.

We computed a “region bias score” to determine whether the components were driven by one region or whether all regions contributed to the component. This was quantified as the square root of the average squared eigenvector elements per region. That produces a 3 element vector, which we normalized to sum to 1. The region bias score was defined as the Euclidean distance between this empirical vector and an “ideal shared region” vector of [1 1 1]/3. The idea is that if all brain regions have average eigenvector components that are equal in magnitude, then that vector will be close to [1 1 1]/3, and thus the empirical distance to the ideal vector will approach zero. As one or two regions start to dominate the component, the normalized average eigenvector elements vector (e.g., producing an empirical vector of [0.6 0.3 0.1]) will move further away from the ideal vector. The maximum possible distance is 1.

Subspace dimensionality was computed via permutation testing. The ability to derive inferential statistical values is one of the important advantages of generalized eigendecomposition over descriptive decompositions such as PCA or ICA. The idea here was to generate a distribution of maximal eigenvalues that could be expected under the null hypothesis that **S** and **R** contain the same information (note from equation 3 that the expected eigenvalue under the null hypothesis is 1, but maximum eigenvalues could be larger due to sampling variability). To generate this empirical null-hypothesis distribution, we randomly assigned each 2-second segment to average into the **S** or **R** covariance matrices. From each iteration, the largest generalized eigenvalue was stored. After 200 iterations for each frequency, the maximum of the largest eigenvalues was taken as the most extreme eigenvalue that can be expected under the null hypothesis that there are no differences between the **S** and **R** matrices. The number of actual eigenvalues (from the analysis without shuffling) above this extreme H0 value was taken as the dimensionality of the subspace. Note that this permutation method accounts for multiple comparisons over M components because it selects the most extreme value of M components on each iteration. Cleaning the covariance matrices via Euclidean distances was performed during permutation testing as described above.

Entropy was computed for each data channel using k=40 bins for discretization.

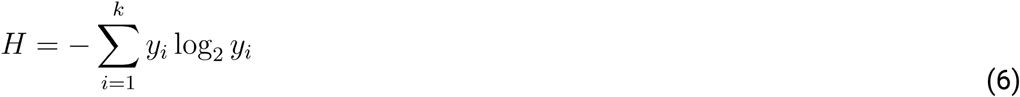

Finally, within-frequency, inter-component phase synchronization was computed via the weighted phase-lag index (Vinck et al., 2011), which is a modification of phase synchronization designed to remove any possible artifacts of volume conduction. This was important for our analyses because all networks were derived by different weightings of the same channels, and because the separate components at the same frequency were not constrained to orthogonality.

## Results

### Data matrices and narrowband source separation

We created channels X time data matrices with 50-80 channels per animal (28-32 LFP channels plus all detected multiunits) (Figure 1), and applied a dimensionality-reduction and guided source-separation method that isolates features of the data that maximally separate narrowband from broadband activity based on generalized eigendecomposition (GED) of covariance matrices (Cohen, 2017). GED was applied after narrowband filtering the data from 2-200 Hz in 100 logarithmically spaced steps, producing a succession of narrowband components. Each component is a weighted average of channels that maximizes energy at that frequency. There are multiple components per frequency that were sorted according to their eigenvalue, which encodes the separability between the narrowband and broadband energy.

Figure 2 illustrates results from one example recording session. This example highlights several consistent features that are expanded on later, including (1) different frequencies engage different electrodes across different regions; (2) some frequencies (e.g., theta) recruit multi-regional networks whereas other frequencies preferentially engage one or two regions; (3) large-scale networks were dominated by LFP whereas multiunits made relatively little (though significant) contributions; (4) the local regional referencing ensured that the components reflected the coordination of multiple local dipoles (seen as the balance between blue and red colors in the map) instead of long-range volume-conducted fields. The components time series had non-Gaussian distributions, indicative of true signals rather than noise, which is expected to be Gaussian-distributed (Figure 2-1)

**Figure 2.**
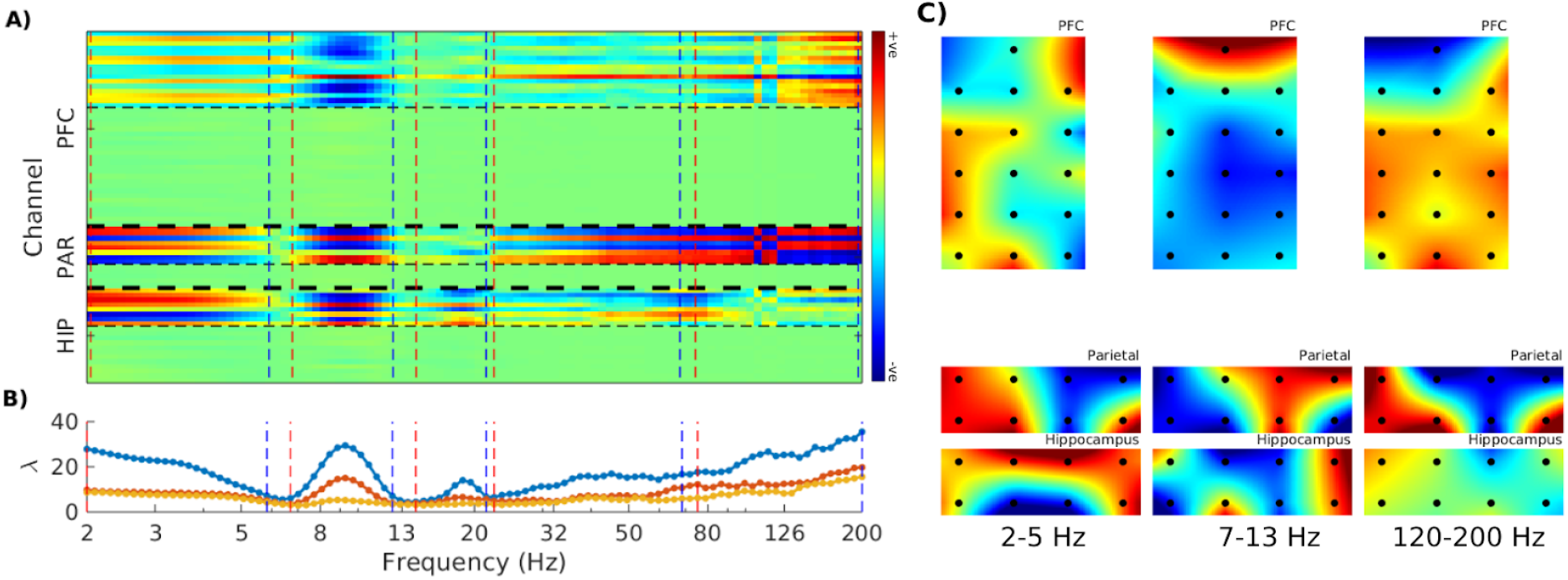
Generalized Eigendecomposition enables spectrally resolved source separation across 3 areas for a single recording. A) Spatial maps over all three regions per frequency (each column corresponds to one frequency). The thick horizontal dashed lines show inter-regional boundaries, and thin horizontal dashed lines show within-region boundaries between LFP (top) and multiunit (bottom) channels. Within-region rows are ordered according to the channel index in the dataset, not according to anatomical location. The colors indicate the strength of the contribution of that channel to the brain-wide component (data were per-frequency normalized, so the color values are arbitrary but comparable across frequencies), vertical dashed lines show the empirically defined frequency boundaries (detailed later): red lines indicate the lower bounds of the frequency band and blue lines indicate the upper bounds. B) Eigenspectra from the largest three components per frequency, which highlights that there can be multiple separable components at the same frequency. The map in panel A is only for the top eigenspectrum (blue line). C) Example topographical maps of the anatomical distribution of the filter projections for the indicated frequency ranges. Each black dot is the location of an electrode. In all columns, medial is to the left and anterior is to the top.

### Empirically derived frequency bands

Electrophysiology data are often grouped into frequency bands according to integer boundaries (e.g., 4-10 Hz), which may miss, artificially separate, or artificially combine the rhythms naturally occurring in the brain. We therefore applied a recently established method (gedBounds) to derive empirical frequency bands based on the definition of a “frequency band” as a range of frequencies that have highly correlated spatiotemporal dynamics (Cohen, 2020). GedBounds works by clustering the matrix of squared correlations across the eigenvectors from all frequencies (Figure 3a). It is a purely data-driven alternative to labeling frequencies based on a priori expectations.

**Figure 3.**
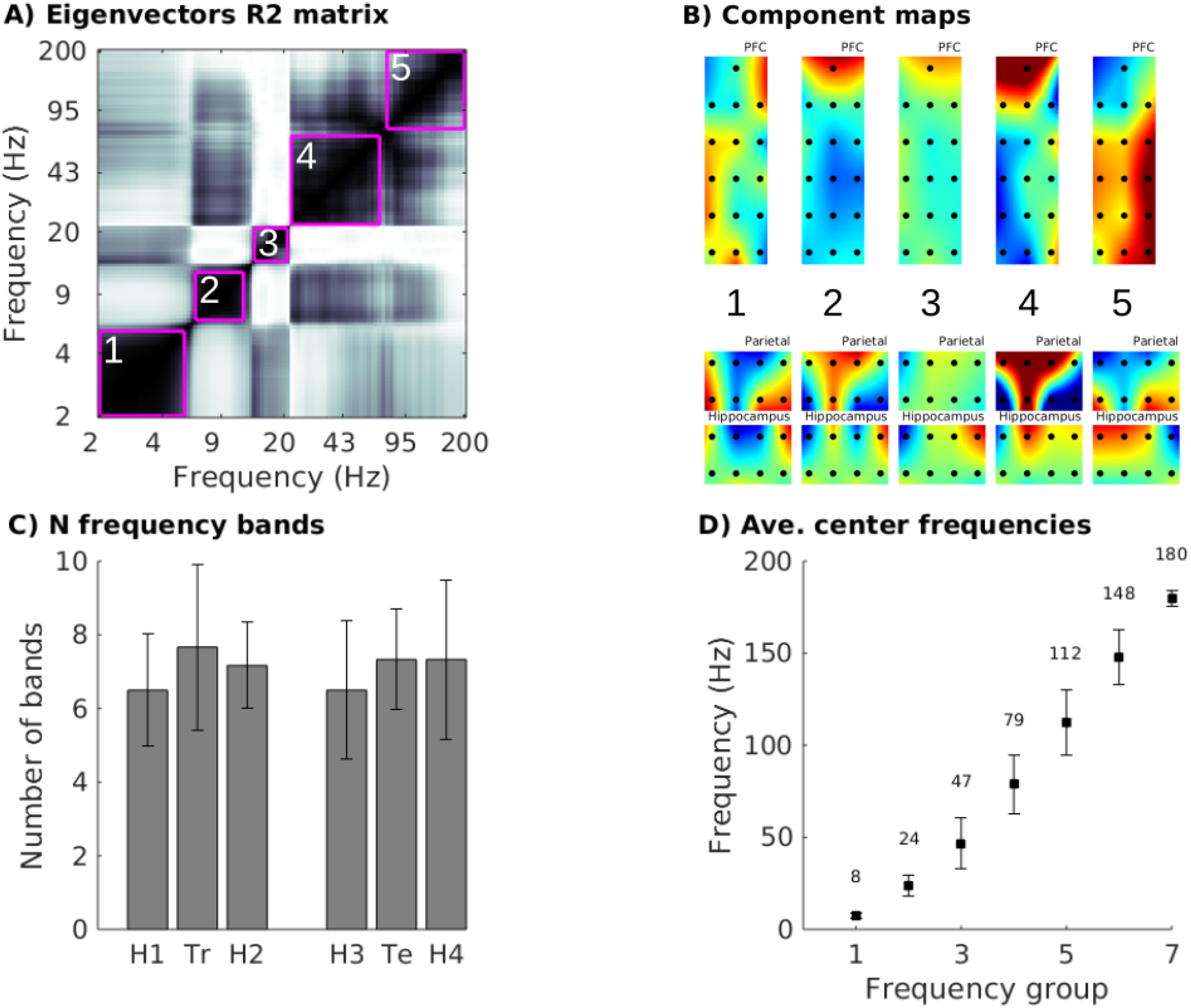
Distinct frequency bands separate clearly in the LFP data with specific spectrotemporal profiles A) R^2^ correlation matrix across all pairs of frequency-specific eigenvectors, with pink boxes drawn around empirically derived clusters (based on the dbscan algorithm), from one recording. The cluster boundaries separate spatially distinct topographies across different frequency ranges. B) Topographical maps of the spatial filter from the frequency bands in panel A. White/black numbers indicate corresponding bands/maps. C) Aggregated results of the number of empirical frequency bands per experiment session (H1-4 indicate home sessions; Tr indicates training session; Te indicates test session). Error bars show standard deviations across the six animals. C) Center frequencies for each group as defined by k-means clustering analysis over animals. Error bars show standard deviations across 100 repeats of the k-means clustering algorithm with different random initializations, and the numbers above each data point shows the average center frequency from that band.

This analysis revealed an average of 7 bands in the range of 2-200 Hz (Figure 3b). The number of frequency bands was not significantly different between experiment sessions (one-way ANOVA: F(5,25)=.45). Average center frequencies were computed by k-means clustering on the empirical frequencies. Because k-means can produce different clusters on each run, we re-seeded the clustering 100 times. The average cluster center frequencies, along with their standard deviations, are shown in Figure 3c.

These results show that grouping electrophysiology time series into spectral bands has an empirical basis and is not arbitrary or an artifact imposed by narrowband filtering. The empirically derived frequency ranges varied over animals and task sessions, and were not systematically affected by the task session. However, we treated frequency as a continuous variable in subsequent analyses rather than grouping into discrete bins.

### Component reproducibility

The anatomical targets of the electrode implants were identical in all animals. However, individual variability in functional organization can mean that the GED patterns are idiosyncratic and thus different across animals. Likewise, if the spatiotemporal patterns that GED isolates reflect stable features of the brain, then the patterns should be highly similar in different experiment sessions within the same animal. On the other hand, it is possible that the spatiotemporal patterns are dynamic and are more affected by cognitive factors than by individual differences.

To address questions about component map reliability, we measured map reproducibility, quantified as spatial correlations, both across experiment sessions within each animal, and in the same session across animals. When pooling across all experiment sessions, we observed robust within-animal component topographies (R^2^ spatial correlations in the range of 0.4 to 0.8 over the frequency spectrum; see Figure 4a). In contrast, spatial correlations across animals were low, with averaged R^2^ values below 0.2. Because the decompositions were performed on the data from each session independently, this pattern of results indicates that (1) the components were stable within each animal over different sessions (over the course of the ~2-hour recording), and that (2) component maps are idiosyncratic, with different spatial patterns in different animals.

**Figure 4.**
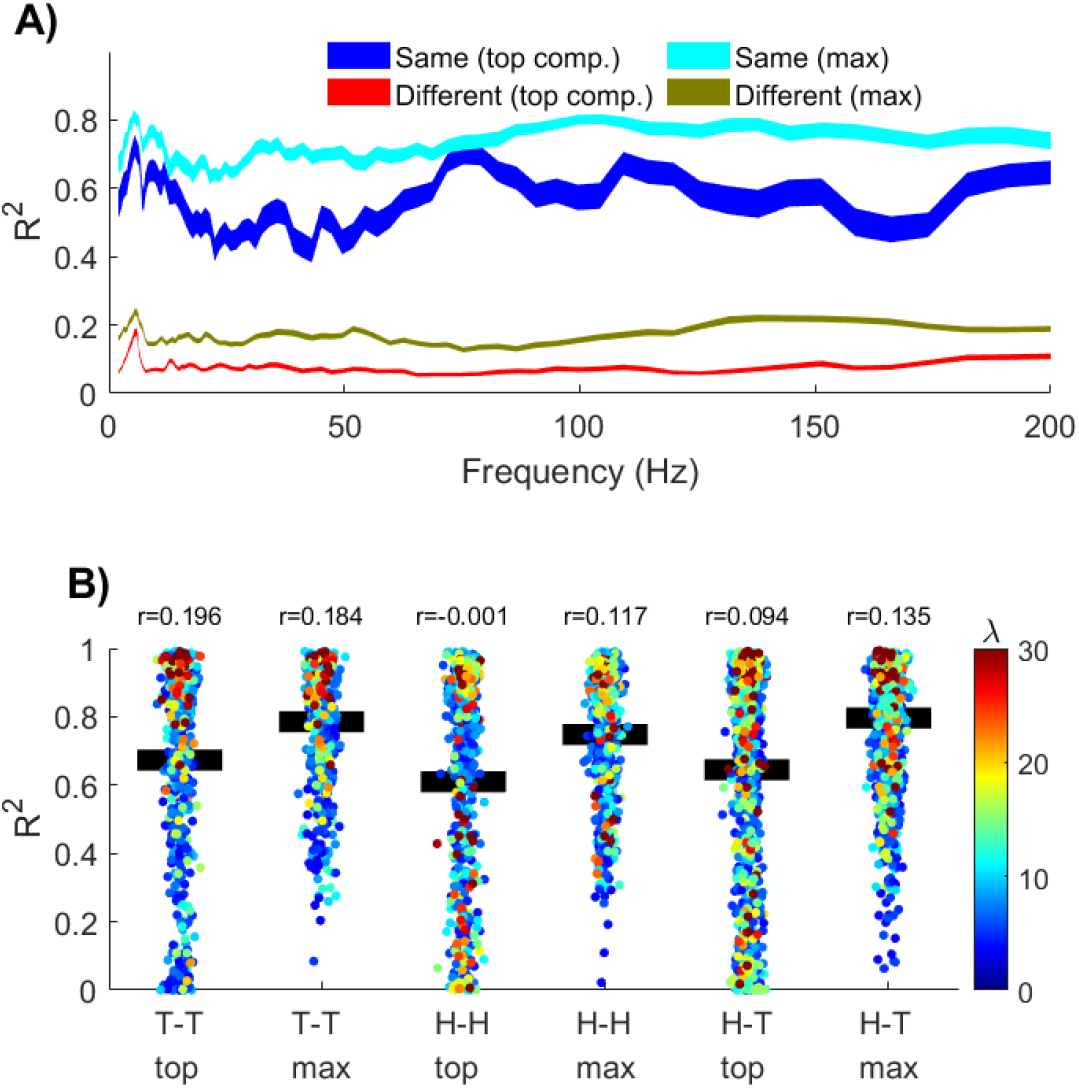
Component topographies are reproducible within animals in different sessions, yet differ across animals. A) R^2^ spatial correlations per frequency. The analysis was run on the components with the largest eigenvalue per frequency (“top comp.”), and by selecting the largest correlation amongst the top two components (“max”). B) Each individual correlation, separated according to the experiment sessions from which the spatial map pairs were drawn (“T-T” indicates train-test pairs, “H-H” indicates home-home pairs). Black bars indicate the mean R^2^. The color of each dot is the average of the eigenvalues of the component pair (which indicates the separability of the narrowband from broadband signals), and the r-value on top of each column is the correlation between the spatial map R^2^ and the average eigenvalue.

The spatial correlations described above were done using only the component with the largest eigenvalue for each session and each frequency. It is possible that the same neurophysiological network was identified as “component 1” in one experiment session and “component 2” in a different session. We therefore modified the correlation analysis to compute the four unique correlations across the top two components from each session/frequency, and stored only the largest correlation coefficient. Although this selection procedure is biased because we selected the strongest correlation out of a set, the same bias was applied within- and across-animals. The correlations were overall stronger, but the conclusion is the same as when correlating only the top components: spatiotemporal patterns were stable within animals, and variable across animals.

We next assessed whether the maps were modulated by the different experiment sessions by separating R^2^ values according to experiment session. The scatter plots in Figure 4B show all frequencies (each dot is an animal-frequency pair), but we averaged frequencies together for the statistics because Figure 4A indicates comparable relationships across the frequency domain. We then tested the correlation coefficients in a one-way ANOVA with the factors train-test, home-home, and train/test-home. In other words, we tested whether the maps were more similar to each other when the animals were in a similar experiment context. However, this effect was not statistically significant (F(2,10)=2.17, p=.16).

Inspection of the distribution of R^2^ values in Figure 4b show considerable spread of the correlations, which was only partially resolved by selecting the maximum correlation of the top two components. We suspected that at least some of this variation could be due to the separability of the components from broadband. “Separability” in a GED analysis is quantified as the eigenvalue, which is the multivariate ratio between the narrowband from the broadband covariance matrices along the direction of the eigenvector. We therefore correlated the R^2^ values with the average of the eigenvalues of each component-pair. Most correlations between map-similarity and eigenvalue were in the range of 0.1-0.2. Thus, it appears that — to some extent — the narrowband components that are better separated from the background spectrum are more likely to be stable over time.

### Region-specificity of components

Given that our data matrices included signals from three brain regions, we next determined whether the components truly reflected inter-regional temporally coherent networks, or whether they were driven by a single region. This was assessed through a regional bias score, in which a score of zero indicates exactly equal contributions from all three regions, whereas a bias score of one indicates that the component is driven entirely by one region with no contributions from the other two regions (Figure 5A).

**Figure 5.**
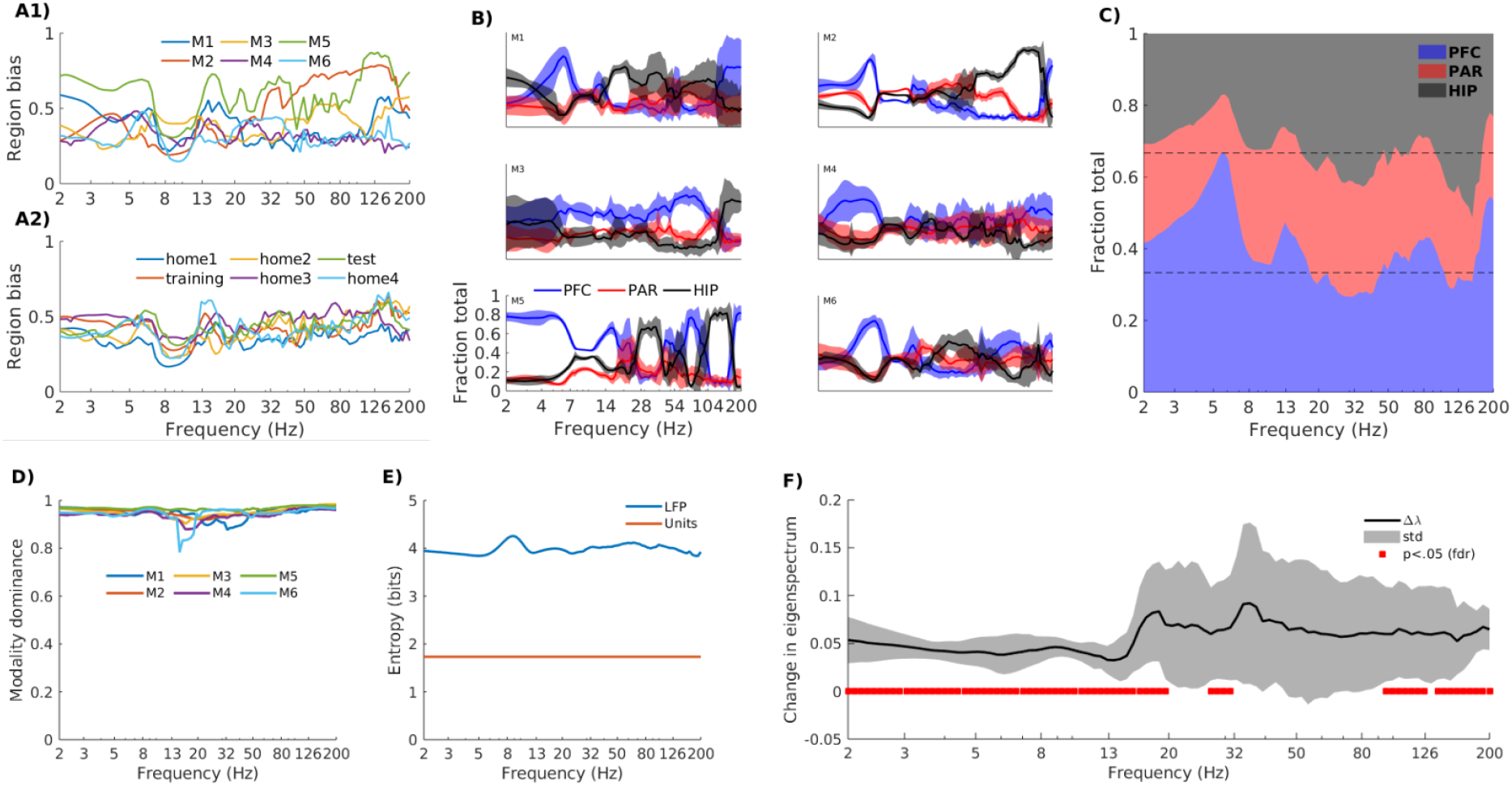
All recorded regions contributed to the components per frequency, with some frequencies showing regional dominance. A) The region bias index for each animal (A1) and averaged over animals for each experiment session (A2). Values close to 0 indicate equal spread of components across all three brain regions, whereas values close to 1 indicate that a single region dominates the component. B) The fraction of total component energy attributable to each region, normalized to the sum over all three regions (thus, the sum per frequency is 1). Each panel is a different animal, averaged over experiment sessions. Patches indicate one standard deviation above and below the mean across sessions, which illustrates the reproducibility of these characteristics over time (six sessions spanning 2 hours). All panels have the same tick marks and axis labels as the lower-left panel. The group-average regional fractions are shown in panel C. Horizontal lines at 1/3 and 2/3 indicate equal contribution of all three regions to the component. D) The modality dominance spectrum quantitatively showed that components were predominantly driven by LFP instead of by multiunits. B) Entropy spectrum shows that LFP channels had higher entropy compared to the multiunits (multiunits’ entropy is the same for all frequencies). F) The multiunits made significant contributions to the components over most frequencies except in the range of 20-90 Hz. Positive values indicate better separability when multiunits are included. The black line is the average over all animals, and the surrounding patch indicates one standard deviation around that average. Red lines show significant changes relative to zero at p<.05, FDR corrected for multiple comparisons over frequencies.

The bias scores were mostly between 0.4 and 0.6 within each animal (Figure 5A), indicating that all three regions contributed to the components to varying degrees. The frequency range that stood out was theta, which exhibited a notable dip in the bias score. Thus, all three brain regions contributed to large-scale networks in the theta range.

This bias score is an aggregate measure; we next investigated the contributions of each region to each frequency, separately for each animal. Figure 5B shows both diversity and commonalities in the regional contributions across the different animals. In these plots, overlapping lines at y=1/3 indicates that all three regions contributed equally to the components, whereas regional dominance is reflected by a separation of lines on the y-axis. Figure 5C illustrates the commonalities across all six animals that are identified through averaging. For example, across animals, PFC generally dominated the low-frequency (<8 Hz) networks whereas the hippocampus generally dominated high-frequency networks between 80-150 Hz.

#### Contributions of LFP vs. multiunits

We next investigated the relative contribution of spikes and LFPs to the components. This was quantified as modality dominance (Zuure et al., 2020), which is the normalized difference between the root-mean-square of the LFP eigenvector elements and the root-mean-square of the multiunit eigenvector elements. A modality dominance value of zero indicates equal contribution of LFP and multiunits, whereas a value of one indicates no contribution of multiunits (a value of minus one would indicate no contribution of LFP channels).

The modality dominance values were close to one for all animals, recording sessions, and frequencies (Figure 5D). This was not attributable to a difference in signal scaling between LFP and multiunits, because all time series signals were normalized to a mean of zero and a variance of one. However, normalizing to the first and second statistical moments does not preclude the possibility of differences in higher-order statistical characteristics. For example, the LFP channels had overall higher entropy (around 4 bits, averaged over all channels, animals, and experiment sessions) compared to the multiunits (1.7 bits on average) (Figure 5E).

On the other hand, it was not the case that multiunits made no contributions to the GED-identified networks. We re-ran the source separation for each frequency, excluding all multiunits from the dataset, and computed a t-test at each frequency between the top eigenvalues from the multiunit-including and multiunit-excluding datasets. The difference was statistically significant (correcting for multiple comparisons using the false discovery rate method (Benjamini and Hochberg, 1995) for most frequencies except around 30-90 Hz (Figure 5F).

Thus, the (Gaussian-smoothed) multiunits made a minor though statistically significant contribution to the matrix decomposition. This overall pattern is not surprising, considering that the LFP samples a larger volume and thus more neurons. On the other hand, there were more multiunit channels in the data matrix than LFP channels, and many of our multiunits may have reflected a combination of several neurons; thus, we interpret this finding to indicate that LFP signals are a richer source of information regarding cross-regional network formation than are action potentials.

### Within-frequency component dimensionality

The eigenvectors from the GED analysis carve out a low-dimensional subspace of narrowband activity, and we defined the dimensionality of that subspace as the number of eigenvalues that were larger than a significance threshold based on a null-hypothesis distribution of eigenvalues derived from permutation testing (Zuure et al., 2020).

The subspace dimensionality ranged from 2 to 16, and generally increased with higher frequencies (Figure 6A-B). Higher dimensionality corresponds to the number of statistically separable networks operating at the same frequency. It is noteworthy that there is no pronounced “bump” in the theta range (~4-10 Hz).

**Figure 6.**
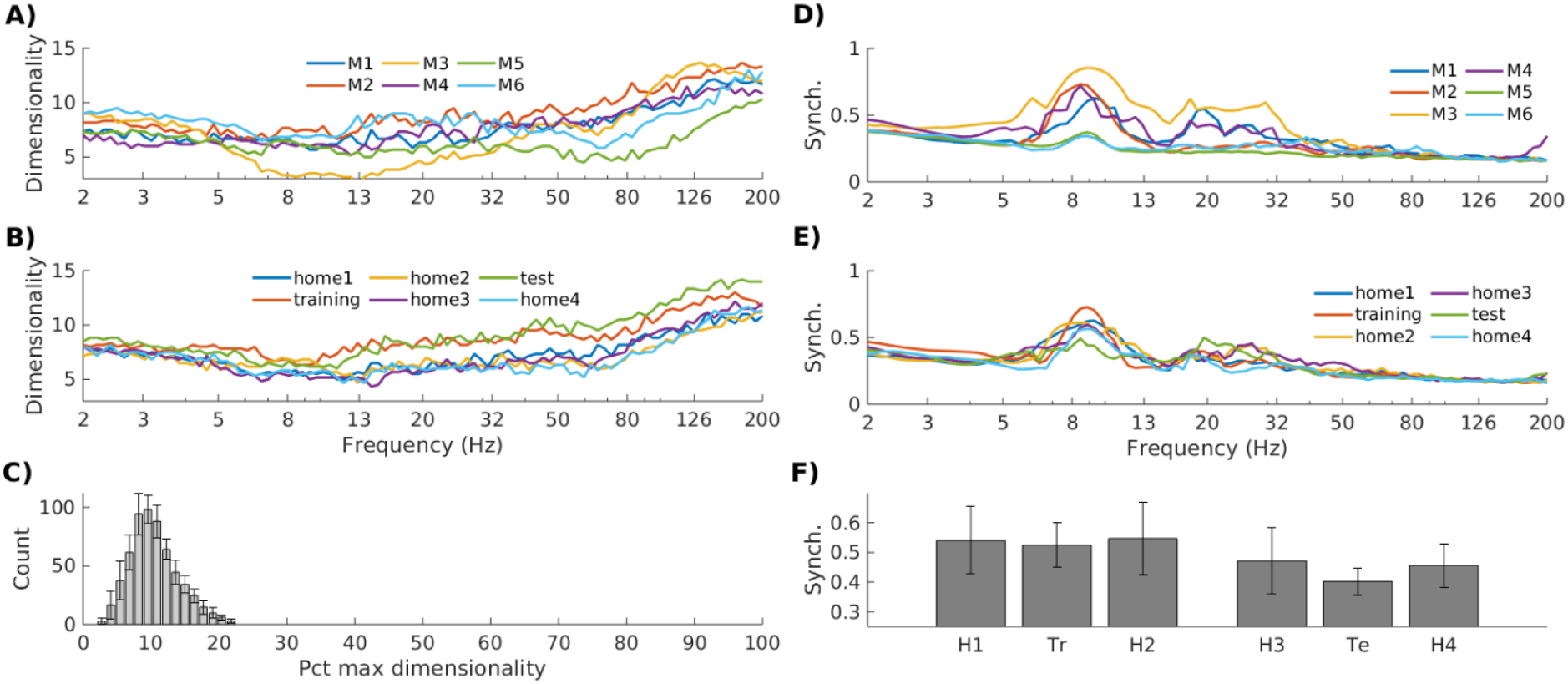
Generalized eigendecomposition reveals that narrowband subspaces are multidimensional (quantified as the number of statistically significant components), and components within each frequency are partially synchronized but non-redundant. A-B) Subspace dimensionality across animals (A) and experiment sessions (B). C) The distribution of all component dimensionalities, normalized to percent of the maximum possible dimensionality (the rank of the covariance matrices), revealed that the narrowband components spanned around 10% of the total possible signal dimensionality. D-F) Phase synchronization between the top two components per frequency indicates both coordination and independence across within-frequency networks. Volume-conduction-independent phase synchronization tended to decline with frequency except for a prominent peak in theta/alpha (~7-13 Hz) and a smaller prominence in beta (~15-30 Hz). The patterns were similar over different animals (D) and different sessions (E). F) Average synchronization in the theta/alpha range for the different sessions.

Note that this measure is not the total dimensionality of the signal; it is the dimensionality of the subspace that differentiates narrowband from broadband activity. Normalizing these raw numbers to the total dimensionality of the signal (assessed as the rank of the corresponding data covariance matrix) revealed that most narrowband subspaces occupied around 8-10% of the total signal space (Figure 7C).

We investigated the dynamics within these subspaces by computing a volume-conduction-independent measure of phase synchronization (weighted phase lag index) between the top two components for each frequency and task session (note that GED eigenvectors are not constrained to orthogonality as with PCA, and thus within-frequency components can be arbitrarily strongly correlated as long as they remain linearly separable). Synchronization strength varied between around .2 and .6 depending on the frequency, with strongest synchronization around theta and a smaller departure from the 1/f decay around the beta band (Figure 6D-F). A repeated-measures ANOVA on session differences in the 7-12 Hz range indicated no main effect of task session (F(5,25)=1.3, p=.29).

## Discussion

In this study, we explored multivariate LFP and multiunit data from three brain regions in awake behaving mice using a combination of established and novel multivariate analysis methods to decompose the data into multiple spatial-spectral-temporal modes. We found that these were stable within each animal, but variable across animals. These findings reveal a rich and multidimensional landscape of brain dynamics that highlight the complexity of on-going neural activity.

### Feature-guided source separation identifies large-scale narrowband networks

There are several dimension-reduction methods that are regularly applied in neuroscience, including principal and independent components analyses, factor analyses, and Tucker decompositions (Cunningham and Yu, 2014). It is often unclear which algorithms or which parameters are optimal (Cohen and Gulbinaite, 2014), and different algorithms can give similar or divergent results (Cohen, 2017; Delorme et al., 2012) depending on their maximization objectives.

GED has several advantages, including that it (1) separates narrowband from broadband activity while holding constant behavioral, cognitive, and other factors; (2) reduces the impact of artifacts or non-brain sources that have a relatively wide frequency distribution; (3) is amenable to inferential statistical thresholding, whereas other decompositions are descriptive and thus selecting components for subsequent interrogation may be subjective or biased; (4) takes into account both spatial and temporal dynamics instead of only spatial or only temporal features; (5) has higher signal-to-noise ratio characteristics and is more accurate at recovering ground truth simulations compared to principal or independent components analyses (Cohen, 2017; de Cheveigné and Parra, 2014; Nikulin et al., 2011; Zuure and Cohen, 2020).

An important finding here is the discovery that a single frequency band can group multiple distinct but spatially overlapping networks. In typical univariate or bivariate analyses, the LFP from a single electrode is treated as an independent statistical unit, based on the implicit assumption that the volume of tissue recorded by an electrode contains only one functional circuit. But a more likely scenario is that each electrode records a mixture of signals from multiple local circuits in the scale of hundreds of microns to a few mm, particularly in the presence of local coherence (Lindén et al., 2011). Thus, LFP is prone to the same kind of source mixing that affects MEG and EEG (Nunez and Srinivasan, 2006), though to a lesser extent. This, however, is fortuitous for multichannel recordings, because it means that linear separation methods that have been established in the EEG community are likely to be fruitful in invasive recordings.

The high reproducibility across sessions within each animal (Morrow et al., 2020), coupled with the low reproducibility across animals, suggests that the large-scale networks that manifest as coordinated LFP dynamics develop in idiosyncratic ways across different individuals. This result, of course, does not invalidate the standard neuroscience approach of targeting the same XYZ coordinates in different individuals and pooling the results together, but our findings highlight that the spatial topographies of larger networks may be unique across individuals, which should be taken into consideration in future studies.

### The special role of theta in large-scale network formation

The theta frequency band, typically defined as 4-10 Hz in rodents in 4-8 Hz in humans, is widely implicated in a large range of cognitive processes, including spatial exploration, memory, motor function, and executive functioning. Clearly, there is no simple mapping of frequency band to cognitive process and indeed, even the same brain regions can generate multiple sources of theta independently (López-Madrona et al., 2020; Zuure et al., 2020), which may serve different cognitive functions (Mikulovic et al., 2018; Töllner et al., 2017). In the rodent brain, theta is most robust in the hippocampus, but also synchronizes with independent theta generators in the medial prefrontal cortex (O’Neill et al., 2013; Sigurdsson and Duvarci, 2015). Intracranial EEG studies in humans have confirmed that theta synchronization is widespread and linked to cognitive operations (Solomon et al., 2017).

The theta band stood out in many of our analyses, for example by having relatively strong within-frequency, cross-component synchronization (Figure 6), sub-Gaussian kurtosis (Figure 2-1), and roughly equal contribution from all three regions (Figure 5). Additionally, theta-band networks appeared to have the most anatomically consistent topographies across animals (see the small peak around theta in Figure 4a). On the other hand, the subspace dimensionality of theta was not higher than other frequencies (Figure 6a-b), suggesting that the theta is important for computational reasons, and is not simply the dominant frequency in general.

### LFP vs. multi-unit contributions to large-scale networks

It is perhaps unsurprising that the multiunits made relatively little statistical contribution to the narrowband components, considering that LFP samples a larger volume, has more signal complexity, and can be meaningfully separated into narrow frequency bands. On the other hand, the multiunits were recorded from the same electrodes, added unique information to the narrowband covariance matrices, and improved the overall separability of the narrowband components from broadband across most frequency ranges.

It is possible that LFP carries most of the inter-regional signaling (Yuste, 2015), considering that LFP reflects a multitude of intra- and extracellular processes (Buzsáki et al., 2012; Reimann et al., 2013) that are modulated by population dynamics of excitatory and inhibitory cells (Mitzdorf, 1985). It is also possible that spikes carry important information that is spatiotemporally targeted and sparse, and therefore make contributions at a spatial scale smaller than what we investigated. Indeed, the eigendecomposition will prefer larger patterns of covariance over patterns driven by a single data channel. On the other hand, LFP is generally considered a proxy of the local input to a circuit while spikes are considered a proxy of the output of the circuit. Nonetheless, multiunits and LFP are rarely incorporated into the same data matrix as we have done, so their relative contributions should be quantitatively evaluated rather than intuitively inferred.

### Implications for novelty and memory

The main network characteristics we identified were not significantly different across the task sessions. This seems to suggest that these network dynamics reflect stable neural architectures as opposed to fluctuating cognitive states.

It is, however, possible that behavior modulates these network dynamics at a faster timescale than experiment sessions. Indeed, neural signatures of novelty processing may be transient, lasting only hundreds of ms (Ranganath and Rainer, 2003) or tens of seconds when first introduced to a novel environment (França et al., 2014). For example, our camera tracking data (not reported here) revealed that animals tended to explore the objects for brief windows of time — sometimes only a few hundred ms. These windows may have been too brief for sufficient neural network estimation, and due to the novelty of the data analysis methods, we chose to focus on characterizing the neural networks using maximal data to ensure high data quality. This could be explored in future studies by ensuring that a particular behavior is expressed for a longer period of time.

## Acknowledgements

We thank Mihaela Gerova for assistance with data cleaning and preparation.

## Extended data

**Figure 1-1.**
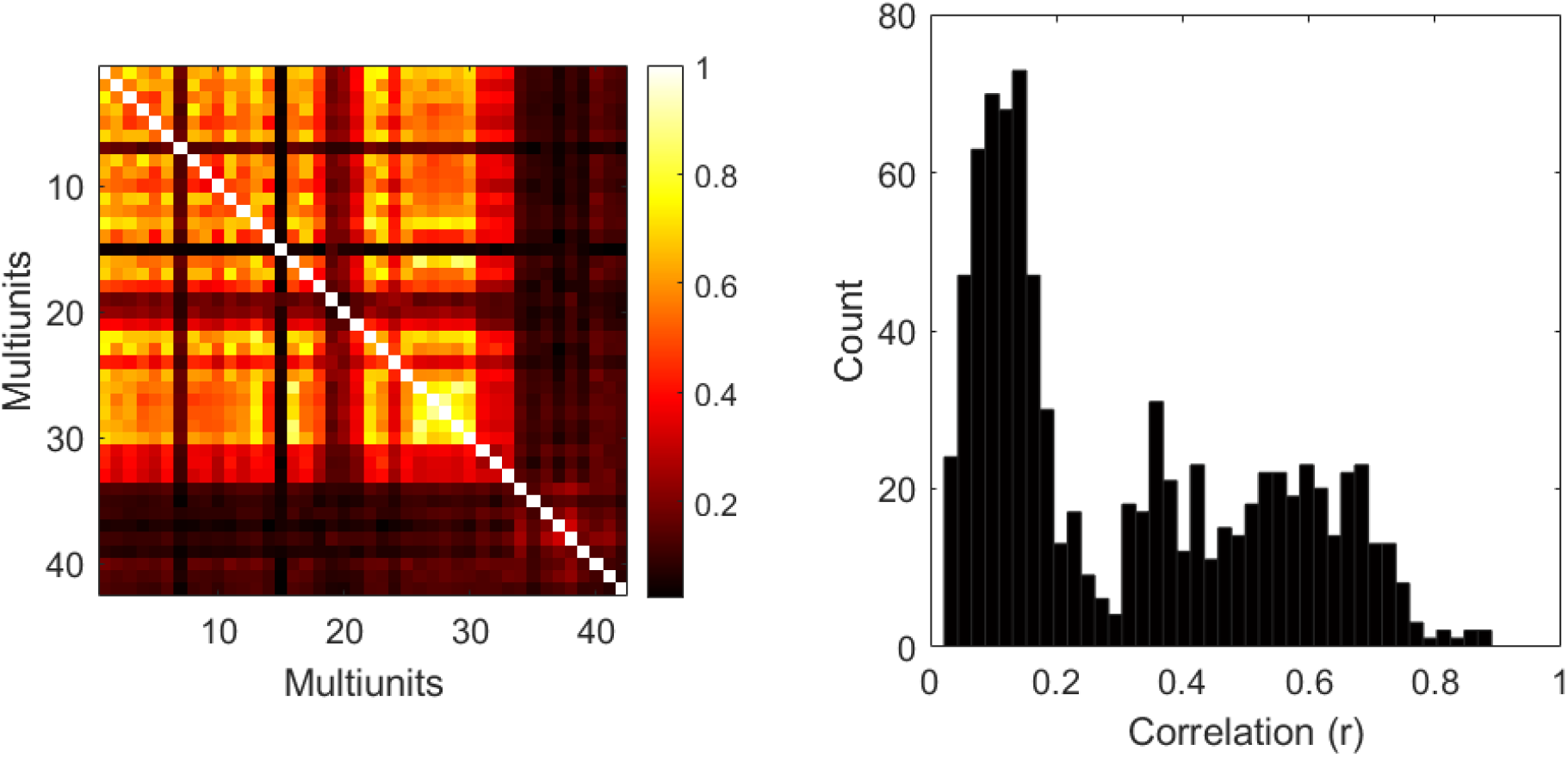
Left plot shows an example multiunit correlation matrix from one recording session. The right plot shows a histogram of all unique off-diagonal correlation values. These plots illustrate that our spike-sorting approach was not overly contaminated by identifying the same units on multiple channels.

**Figure 2-1.**
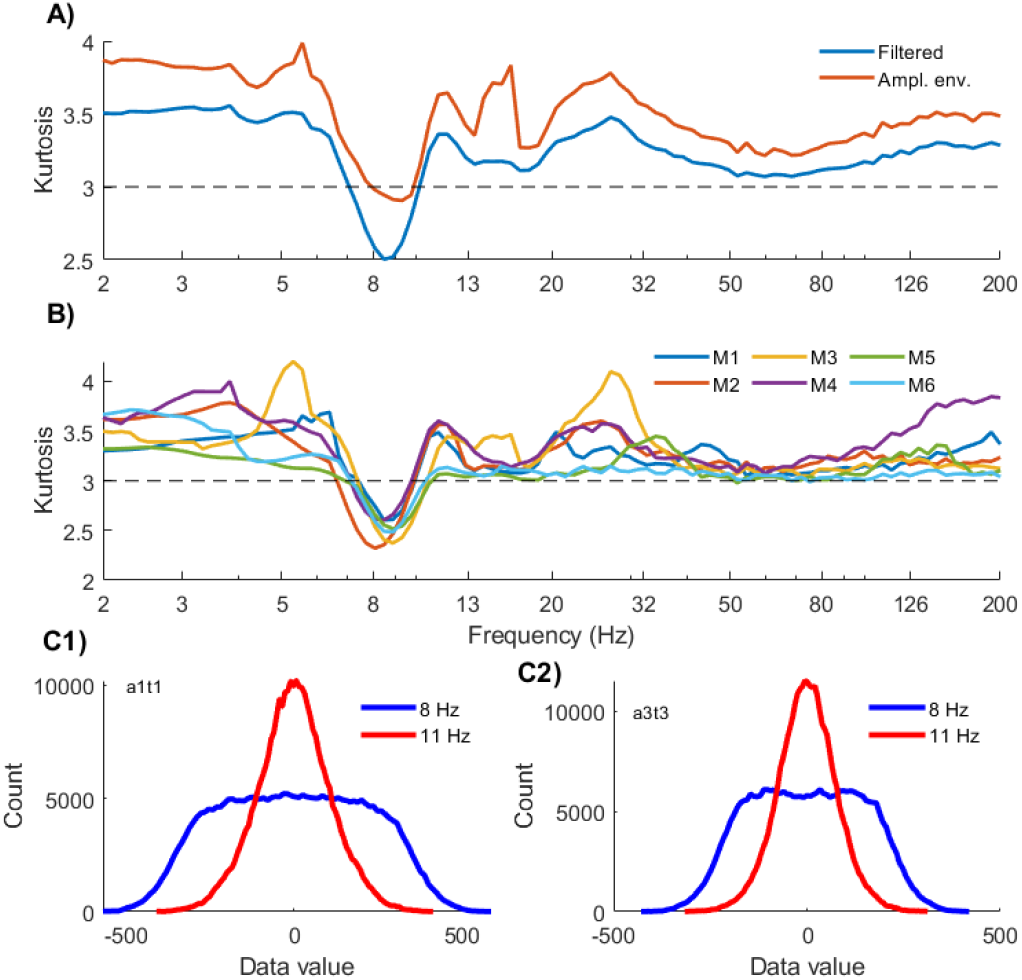
Kurtosis, a measure of non-Gaussianity of a distribution (see text below), computed on frequency-specific component time series. The red and blue lines in panel A show kurtosis per frequency for the narrowband-filtered time series (blue) and amplitude envelope (red), averaged over all animals and sessions. The horizontal dashed line indicates the expected kurtosis of a pure Gaussian distribution. B) Kurtosis over frequencies for each animal separately. Note the striking decrease in kurtosis in the theta band in all animals. C) Example time series histograms illustrating the platykurtic effect at 8 Hz and 11 Hz for two different animals and sessions.

### Distribution shape via kurtosis

Non-Gaussianity is considered an indicator of an information-rich signal. This comes from the central limit theorem, which leads to the assumption that random noise, and random linear mixtures of signals, will produce Gaussian distributions. We therefore quantified the kurtosis (4th statistical moment of a distribution; the kurtosis of a pure Gaussian distribution is 3) as a measure of the non-Gaussianity of the component time series. We computed kurtosis for the narrowband filtered signal and its amplitude envelope at each component.

Component time series kurtosis was computed as the 4th statistical moment of the component time series. We extracted kurtosis from both the real part of the narrowband signal and the amplitude envelope (extracted via the Hilbert transform). The amplitude envelope had overall higher kurtosis (Figure 2-1), which is not surprising considering that amplitude is a strictly non-negative quantity.

Nearly all frequencies had kurtosis higher than 3, indicating leptokurtic distributions characterized by narrow peaks and fatter tails. This is consistent with suggestions that brain activity is characterized by extreme events and long-tailed distributions (Buzsáki and Mizuseki, 2014). Curiously, all six animals exhibited a dip in kurtosis in the theta band (~9 Hz) (Figure 2-1b), indicating a platykurtic distribution with data values clustered towards zero and relatively fewer data points having extreme values (the tails of the distributions) (Figure 2-1c). This may be related to the known sawtooth-like shape of hippocampal theta (Scheffer-Teixeira and Tort, 2016).

Note that unlike independent components analysis, GED is based purely on the signal covariance (second moment) and not on any higher-order statistical moments. Thus, non-Gaussian distributions are not trivially imposed by the decomposition method, but instead arose from the data without bias or selection.

## References

Alishbayli A, Tichelaar JG, Gorska U, Cohen MX, Englitz B (2019) The asynchronous state’s relation to large-scale potentials in cortex. J Neurophysiol 122:2206–2219.

Benjamini Y, Hochberg Y (1995) Controlling the False Discovery Rate: A Practical and Powerful Approach to Multiple Testing. Journal of the Royal Statistical Society: Series B (Methodological).

Buzsáki G, Anastassiou CA, Koch C (2012) The origin of extracellular fields and currents — EEG, ECoG, LFP and spikes. Nat Rev Neurosci 13:407.

Buzsáki G, Mizuseki K (2014) The log-dynamic brain: how skewed distributions affect network operations. Nat Rev Neurosci 15:264–278.

Carandini M (2005) Do We Know What the Early Visual System Does? Journal of Neuroscience.

Cardoso JF (1999) High-order contrasts for independent component analysis. Neural Comput 11:157–192.

Cohen MR, Kohn A (2011) Measuring and interpreting neuronal correlations. Nat Neurosci 14:811–819.

Cohen MX (2020) A data-driven method to identify frequency boundaries in multichannel electrophysiology data. biorxiv.

Cohen MX (2019) A better way to define and describe Morlet wavelets for time-frequency analysis. Neuroimage 199:81–86.

Cohen MX (2017) Comparison of linear spatial filters for identifying oscillatory activity in multichannel data. J Neurosci Methods 278:1–12.

Cohen MX, Gulbinaite R (2014) Five methodological challenges in cognitive electrophysiology. Neuroimage 85 Pt 2:702–710.

Cunningham JP, Yu BM (2014) Dimensionality reduction for large-scale neural recordings. Nat Neurosci 17:1500–1509.

de Cheveigné A, Parra LC (2014) Joint decorrelation, a versatile tool for multichannel data analysis. Neuroimage 98:487–505.

Delorme A, Makeig S (2004) EEGLAB: an open source toolbox for analysis of single-trial EEG dynamics including independent component analysis. J Neurosci Methods 134:9–21.

Delorme A, Palmer J, Onton J, Oostenveld R, Makeig S (2012) Independent EEG sources are dipolar. PLoS One 7:e30135.

França ASC, do Nascimento GC, Lopes-dos-Santos V, Muratori L, Ribeiro S, Lobão-Soares B, Tort ABL (2014) Beta2 oscillations (23-30 Hz) in the mouse hippocampus during novel object recognition. European Journal of Neuroscience.

Gray CM, König P, Engel AK, Singer W (1989) Oscillatory responses in cat visual cortex exhibit inter-columnar synchronization which reflects global stimulus properties. Nature 338.

Gusnard DA, Raichle ME, Raichle ME (2001) Searching for a baseline: functional imaging and the resting human brain. Nat Rev Neurosci 2:685–694.

Haufe S, Meinecke F, Görgen K, Dähne S, Haynes J-D, Blankertz B, Bießmann F (2014) On the interpretation of weight vectors of linear models in multivariate neuroimaging. NeuroImage.

Hebart MN, Baker CI (2018) Deconstructing multivariate decoding for the study of brain function. NeuroImage 180:4–18.

Hubel DH, Wiesel TN (1959) Receptive fields of single neurones in the cat’s striate cortex. The Journal of Physiology.

Jensen O, Mazaheri A (2010) Shaping functional architecture by oscillatory alpha activity: gating by inhibition. Front Hum Neurosci 4:186.

Kohn A, Coen-Cagli R, Kanitscheider I, Pouget A (2016) Correlations and Neuronal Population Information. Annual Review of Neuroscience.

Kriegeskorte N, Kievit RA (2013) Representational geometry: integrating cognition, computation, and the brain. Trends Cogn Sci 17:401–412.

Lindén H, Tetzlaff T, Potjans TC, Pettersen KH, Grün S, Diesmann M, Einevoll GT (2011) Modeling the spatial reach of the LFP. Neuron 72:859–872.

López-Madrona VJ, Pérez-Montoyo E, Álvarez-Salvado E, Moratal D, Herreras O, Pereda E, Mirasso CR, Canals S (2020) Different theta frameworks coexist in the rat hippocampus and are coordinated during memory-guided and novelty tasks. Elife 9.

Lotte F, Guan C (2011) Regularizing common spatial patterns to improve BCI designs: unified theory and new algorithms. IEEE Trans Biomed Eng 58:355–362.

Mikulovic S, Restrepo CE, Siwani S, Bauer P, Pupe S, Tort ABL, Kullander K, Leão RN (2018) Ventral hippocampal OLM cells control type 2 theta oscillations and response to predator odor. Nat Commun 9:3638.

Mitzdorf U (1985) Current source-density method and application in cat cerebral cortex: investigation of evoked potentials and EEG phenomena. Physiol Rev 65:37–100.

Morrow JK, Cohen MX, Gothard KM (2020) Mesoscopic-scale functional networks in the primate amygdala. Elife 9.

Nikulin V, Nolte G, Curio G (2011) A novel method for reliable and fast extraction of neuronal EEG/MEG oscillations on the basis of spatio-spectral decomposition. Klinische Neurophysiologie.

Nunez PL, Srinivasan R (2006) Electric Fields of the Brain: The Neurophysics of EEG. Oxford University Press, USA.

O’Neill P-K, Gordon JA, Sigurdsson T (2013) Theta oscillations in the medial prefrontal cortex are modulated by spatial working memory and synchronize with the hippocampus through its ventral subregion. J Neurosci 33:14211–14224.

Pang R, Lansdell BJ, Fairhall AL (2016) Dimensionality reduction in neuroscience. Current Biology.

Priesemann V, Wibral M, Valderrama M, Pröpper R, Le Van Quyen M, Geisel T, Triesch J, Nikolić D, Munk MHJ (2014) Spike avalanches in vivo suggest a driven, slightly subcritical brain state. Front Syst Neurosci 8:108.

Ranganath C, Rainer G (2003) Neural mechanisms for detecting and remembering novel events. Nat Rev Neurosci 4:193–202.

Reimann MW, Anastassiou CA, Perin R, Hill SL, Markram H, Koch C (2013) A biophysically detailed model of neocortical local field potentials predicts the critical role of active membrane currents. Neuron 79.

Richter CG, Babo-Rebelo M, Schwartz D, Tallon-Baudry C (2017) Phase-amplitude coupling at the organism level: The amplitude of spontaneous alpha rhythm fluctuations varies with the phase of the infra-slow gastric basal rhythm. Neuroimage 146:951–958.

Ritchie JB, Kaplan DM, Klein C (2019) Decoding the Brain: Neural Representation and the Limits of Multivariate Pattern Analysis in Cognitive Neuroscience. Br J Philos Sci 70:581–607.

Scheffer-Teixeira R, Tort AB (2016) On cross-frequency phase-phase coupling between theta and gamma oscillations in the hippocampus. Elife 5.

Sigurdsson T, Duvarci S (2015) Hippocampal-Prefrontal Interactions in Cognition, Behavior and Psychiatric Disease. Front Syst Neurosci 9:190.

Singer W (2009) Distributed processing and temporal codes in neuronal networks. Cogn Neurodyn 3.

Solomon EA, Kragel JE, Sperling MR, Sharan A, Worrell G, Kucewicz M, Inman CS, Lega B, Davis KA, Stein JM, Jobst BC, Zaghloul KA, Sheth SA, Rizzuto DS, Kahana MJ (2017) Widespread theta synchrony and high-frequency desynchronization underlies enhanced cognition. Nat Commun 8:1704.

Timofeev I, Bazhenov M (2005) Mechanisms and biological role of thalamocortical oscillations. Trends in chronobiology research 1–47.

Töllner T, Wang Y, Makeig S, Müller HJ, Jung TP, Gramann K (2017) Two Independent Frontal Midline Theta Oscillations during Conflict Detection and Adaptation in a Simon-Type Manual Reaching Task. J Neurosci 37.

Tomé AM (2006) The generalized eigendecomposition approach to the blind source separation problem. Digital Signal Processing.

Trautmann EM, Stavisky SD, Lahiri S, Ames KC, Kaufman MT, O’Shea DJ, Vyas S, Sun X, Ryu SI, Ganguli S, Shenoy KV (2019) Accurate Estimation of Neural Population Dynamics without Spike Sorting. Neuron 103:292–308.e4.

van Hulten J. A. Cohen M. X Fasc (2020) Low-cost and versatile electrodes for extracellular chronic recordings in rodents. Heliyon 6:e04867.

Vinck M, Oostenveld R, van Wingerden M, Battaglia F, Pennartz CMA (2011) An improved index of phase-synchronization for electrophysiological data in the presence of volume-conduction, noise and sample-size bias. Neuroimage 55:1548–1565.

Wang X-J (2010) Neurophysiological and computational principles of cortical rhythms in cognition. Physiol Rev 90:1195–1268.

Whittingstall K Logothetis (2009) Frequency-Band Coupling in Surface EEG Reflects Spiking Activity in Monkey Visual Cortex. Neuron 64:281–289.

Williamson RC, Doiron B, Smith MA, Yu BM (2019) Bridging large-scale neuronal recordings and large-scale network models using dimensionality reduction. Curr Opin Neurobiol 55:40–47.

Yuste R (2015) From the neuron doctrine to neural networks. Nat Rev Neurosci 16:487–497.

Zuure MB, Cohen MX (2020) Narrowband multivariate source separation for semi-blind discovery of experiment contrasts. bioRxiv.

Zuure MB, Hinkley LBN, Tiesinga PHE, Nagarajan SS, Cohen MX (2020) Multiple midfrontal thetas revealed by source separation of simultaneous MEG and EEG. BioRxiv.

